# Reprogramming of lipid secretome in senescent synovial fibroblasts ameliorate osteoarthritis development

**DOI:** 10.1101/2023.04.15.537012

**Authors:** Jiajie Hu, Dongsheng Yu, Dongmei Wu, Jiasheng Wang, Youzhi Cai, Xiaotian Du, Junfen Ji, Hongwei Ouyang

## Abstract

Osteoarthritis (OA) is one of the most common causes of physical disability among older people and its incidence increases with age. Removal of the senescent cells (SNCs) delays OA pathologies, but little is known about the SNCs in OA synovium and their roles in OA pathogenesis. Here, we reported that RCAN1^+^IL1α^+^ double-positive cells represent a senescent subset of synovial fibroblasts which accelerate cartilage degeneration via saturated fatty acids (SFAs). Using single-cell RNA sequencing and synovial organoids, we found that IL1α+ senescent synovial fibroblasts mainly located at lining layer of human OA synovium and promote cartilage degeneration. We performed genome-wide CRISPR/Cas9 screens to identify novel regulators of cell-surface bounded IL1α in senescent cells. RCAN1, which was a hit as positive regulator of IL1α, governs the cellular senescence and pro-degenerative senescence-associated secretory phenotype (SASP). Mechanistically, we found that RCAN1 mediated the secretion of SFAs from senescent synovial fibroblasts, which promote chondrocytes senescence and cartilage matrix degradation. Intra-articular delivery of synovium-targeted anti-Rcan1 siRNA ameliorates the development of post-traumatic OA in mice, while reducing of SNCs accumulation in synovium and increasing cartilage regeneration. Last, co-culture experiments of human OA cartilage explant with synovial organoids confirmed that RCAN1 silencing in human synovial fibroblasts suppressed chondrocyte senescence and cartilage degradation. Thus, our study revealed a pro-degenerative interaction between RCAN1^+^IL1α^+^ senescent synovial fibroblasts and chondrocytes through secreted lipid mediators in OA progression. And targeted RCAN1 inhibition in senescent synovium is a promising approach for restore the joint homeostasis.

## INTRODUCTION

Osteoarthritis (OA), the most common form of arthritis, is a degenerative joint disease which usually lead to total replacement of the affected joint^1,2^. The root cause of this debilitating chronic illness has not been elucidated, and no disease-modifying therapies currently exist. The pathogenesis of OA is multifactorial, involving both systemic and local factors such as age, obesity, and joint instability^2-4^. Recent studies indicate that removal of damaged or senescent cells (SNCs) in knee joints through a transgenic system attenuates disease progression in a posttraumatic OA mouse model, underscoring the causal role of SNCs in OA pathogenesis^5,6^. Intra-articular injection of senolytics, drugs that selectively kill senescent cells, proves to be therapeutic in mouse OA model^7^. These findings have opened the door for the development of agents and strategies to specifically target SNCs for the prevention and treatment of OA.

OA is primarily characterized by the progressive loss of the cartilage matrix but also involves pathological changes in other joint components such as subchondral bone sclerosis, osteophyte formation, and synovial inflammation^1^. Recent studies about SNCs and OA pathogenesis mainly focused on articular surface and chondrocytes^5,8,9^. For example, both preclinical and clinical studies indicate that an increase in the number of senescent cells in articular cartilage is closely linked to OA development, and microsphere cultures of explanted chondrocytes directly showed a deleterious effect of SNCs on cartilage deposition^7^. In contrast, SNCs located in other joint components such as synovium has been noted^7,10^ but their roles in OA pathogenesis have been largely underestimated. Normal synovial tissue comprises two main layers of specialized columnar fibroblast-like synoviocytes (FLSs) that are interspersed with macrophages^11^. OA synovium commonly exhibit synovial hyperplasia, increased collagen deposition, multiple inflammatory cell infiltrates and presence of sensory neurons^12^. OA-FLS could contribute to OA pathogenesis through: (1) producing inflammatory mediators, (2) destroying cartilage ECM, and (3) activating into myofibroblast-like cells. Generally, fibroblasts are considered to have little heterogeneity. Recently, anatomically discrete, functionally distinct fibroblast subsets with non-overlapping functions have been identified in rheumatoid arthritis (RA) with single cell sequencing^13^. However, it remains largely unknown whether FLS are also phenotypically and functionally heterogenous in OA.

SNCs contribute to disease development through the secretion of proinflammatory cytokines and proteases, as IL1α, IL1β, IL6, IL8, and matrix metalloproteinases (MMP1 and MMP3), referred to as senescence-associated secretory phenotype (SASP) factors^14^. Microsphere cultures of explanted chondrocytes directly showed a deleterious effect of SNCs on cartilage deposition, which suggests that SASP made by SNCs inhibits cartilage regeneration^7^. However, SASP are highly dynamic and have pleiotropic effects on surrounding tissue^15^. Despite the broad biological activities attributed to the SASP, little is known about how it is regulated other than the activation of inflammasome and the classical regulators of inflammation associated with NF-kB activity, such as CCAAT/enhancer-binding protein b (C/EBPb), Mammalian target of rapamycin (mTOR) and p38 mitogen-activated protein kinase (p38 MAPK)^16-18^. In contrast to acute inflammatory responses, the SASP develops relatively slowly, usually manifesting only a few days after the onset of senescent growth arrest; suggesting the existence of a senescence-specific activation mechanism. Recently, accumulating evidence indicates that SASP are sensitive to cellular metabolic states and lipid components of SASP remain understudied ^19-21^. In addition, metabolic disorders are considered to be closely associated with OA, and some aspects of OA are clearly rooted in cell-intrinsic metabolic alterations ^22,23^. However, the cellular source of extracellular metabolites and their roles in OA is not well understood.

The role of ageing joint niche in the pathogenesis of OA has not been fully investigated. Here, combining genome-wide CRISPR/Cas9 knockout screening and single cell analysis, we investigated how senescent synovial fibroblasts contribute to OA pathogenesis and assessed whether the joint homeostasis can be restored by novel anti-senescence approaches. We revealed a degenerative crosstalk between senescent synovial fibroblasts and articular chondrocytes in OA joint mediated by a lipid metabolite-palmitic acid. An OA-associated senescent subpopulation was identified in the synovium, which is RCAN1^+^IL1α^+^ double positive, mainly located at the lining layer of the OA synovium, and exhibit late-stage senescent phenotype and dysregulated lipid metabolism. Finally, we found that RCAN1 promote cellular senescence and saturated fatty acid secretion from senescent synovial fibroblasts and inhibition of RCAN1 in synovium could restore the joint homeostasis in mouse and human models.

## RESULTS

### IL1α^+^ senescent fibroblast accumulates in human OA synovium and promote cartilage degeneration

To define senescent populations and identify genome-wide gene expression patterns, we performed single cell RNA sequencing (scRNA-seq) data of human non-OA (n=3) and OA (n=5, integrated with data from GSE152805) synovium obtained from the patients undergoing knee arthroplasty surgery^24^. After assigning identities to all cell clusters (Fig. 1A, Extended Data FigS. 1), targeted reanalysis of the fibroblast populations on the basis of expression of known fibroblast markers^24,25^ revealed the existence of eight distinct subgroups: 4 lining layer fibroblasts (L1, L2, L3, L4) that express high level of PRG4, CD55 and HTRA1, and 4 sublining layer fibroblasts (S1, S2, S3, S4) that are positive of THY1, IGF1 and CDH11 expression (Fig. 1B-C). To identify the pathway alterations that underlie these fibroblast clusters, we performed gene set variation analysis (GSVA) analysis^26^. Lining fibroblast cluster L2 and sublining cluster S4 marker genes were over-represented in categories related to “replicative senescence”, “stress induced premature senescence” and “positive regulation of IL1A production”, indicating these two clusters represent the “senescent” subpopulation (Fig. 1D). Next, we examined the composition of each cluster in the context of disease and found that the “senescent” lining fibroblast cluster L2 were significantly overrepresented in late-stage OA, while the “senescent” sublining fibroblast cluster S4 does not (Fig. 1E). To corroborate our finding of senescent cells accumulation in OA synovium, we used histological staining of p16 and SA-β-Gal activity, markers that are commonly used to detect senescent cells. Consistent with our scRNA-seq data, we found that senescent cells were particularly abundant in the lining layer of OA synovium (Fig. 1F). Histological staining of PRG4+ lining fibroblasts with IL1α, a pro-inflammatory SASP component, showed percentage of PRG4+ IL1α+ fibroblasts in the synovium was more abundant in OA compared with non-OA synovium (Fig. 1F).

**Fig 1:**
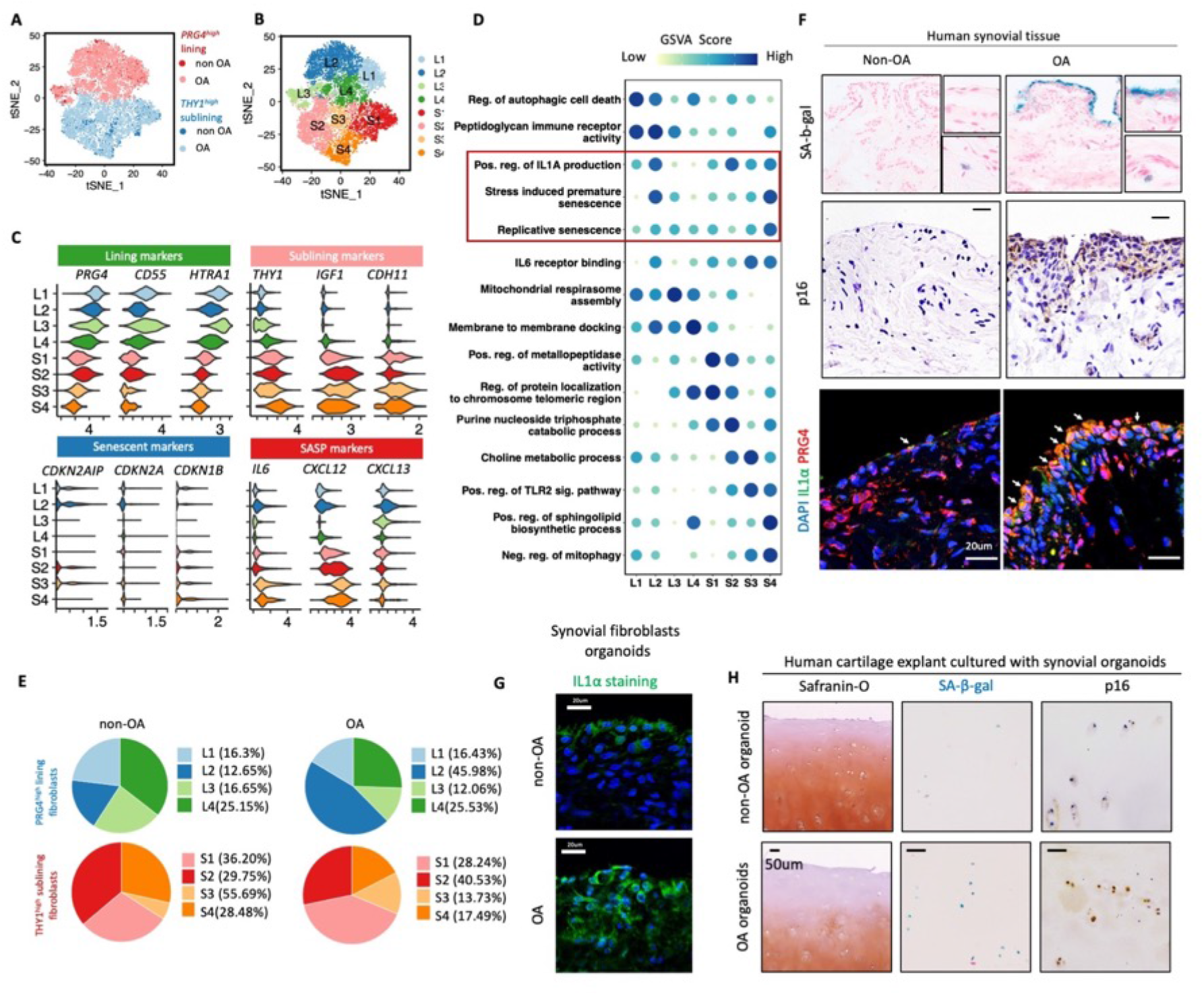
IL1α^+^ senescent fibroblast accumulates in human OA synovium and promote cartilage degeneration. **A**, t-SNE projection showing 8,675 synovial fibroblasts from the synovium of human knee OA joints (n = 5 biological replicates). Cells are colored according to major cell type based on expression of known marker genes. **B**, Unsupervised graph-based clustering of fibroblasts reveals 8 subsets in the human OA joints. **C**, Expression of marker genes (x axes) in the identified OA fibroblast clusters (y axes). The green panels show expression of known lining fibroblast markers. The pink panels show examples of known makers genes of sublining fibroblast. The blue panels show examples of known senescent markers. The red panels show examples of known makers genes of senescent associated secretory phenotype (SASP). **D**, Gene Set Variation Analysis (GSVA)of GO categories suggests different functions for the eight OA fibroblast subsets. **E**, Pie charts (values in percentages; each pie chart represents 100%) showing the composition of each clusters. **F**, SA-β-gal, immunohistological staining of senescent markers (p16 and IL1α) and lining synovial fibroblast marker (PRG4) in non-OA (n=3) and OA (n = 6) synovium. Scale bar, 20μm. **G**, Immunofluorescent staining of IL1α in synovial organoids derived from non-OA and OA synovial fibroblasts. Scale bar, 20μm. **H**, safranin O staining, SA-β-gal and immunohistological staining of p16 in human cartilage explants after coculture with human non-OA or OA organoids for 14 days. Scale bar, 50μm.

It has been noted that the effect of SNCs is pleiotropic, which can be thought of as pro-degenerative or pro-regenerative with regard to the specific context^27,28^. IL1α has been shown to be an upstream regulator of other pro-inflammatory SASP factors during cellular senescence, including IL6, IL8^29,30^. To further verify whether senescent synovial fibroblasts could accelerate the pathological progression of OA, we cocultured human cartilage explants with synovial organoids either from patients with OA or patients without OA. We detected increased expression of IL1α in OA synovial organoids compared with non-OA group (Fig. 1G). Interestingly, we detected increased level p16 and SA-β-Gal activity and decreased level of cartilage extracellular matrix in human cartilage explants after coculture with OA synovial organoids(Fig. 1H). Taken together, we identified a IL1α+ senescent fibroblasts accumulated at lining layer of human OA synovium, which could contribute to chronic synovium inflammation and cartilage degeneration.

### Genome-wide CRISPR knockout screening identified novel regulators for IL1α in senescent cells

To investigate the genetic regulators of IL1α in senescent cells, we performed genome-wide CRISPR/Cas9 knockout library screening in senescent human ESC-MSCs. Human GeCKO v2 CRISPR library A contains 65,386 unique sgRNAs, which targets 19,052 protein-coding genes, and 1864 microRNAs were used to generate a mutant cell pool. ESC-MSCs were infected with the library at a multiplicity of infection (MOI) of 0.3 and selected with puromycin for transduced cells. We then treated the mutant cell pool with doxorubicin for 7 days to establish cellular senescence and cells were sorted based on high or low cell-surface IL1Α expression levels (about 5 million cells/bin) (Fig. 2A). We reasoned that knockout of a IL1α positive regulator gene will decrease cell-surface IL1α level, thus sgRNAs against positive regulator of IL1α would be enriched in low IL1α (“IL1α ^low^”) relative to high IL1α (“IL1^high^”), and that can be determined by high-throughput sequencing. After selection, we achieved about 400× coverage of the library and around 94% of the sgRNA sequences were retained in all samples, which ensure the sufficient read death and library coverage for the CRISPR library screening. From this CRISPR/Cas9 knockout library screening, we identified a subset of sgRNAs targeting 206 genes were significantly enriched (P < 0.05) in the IL1α ^low^ cells when compared to IL1^high^, indicating that these genes might be potential drivers for cell-surface IL1α expression (Fig. 2B). Pathway analysis (DAVID Bioinformatics Resources 6.8) suggested that these genes were involved in UV-damage excision repair, positive regulation of TORC1 signaling and autophagosome maturation, which echoed previous study results (Fig. 2C)^31^. Among the list of genes, we identified previous known SASP regulators, such as BRD4, ATM, HMGB1/2 and GATA4(Fig. 2D). However, most identified genes had not been previously implicated in IL1α or SASP regulation.

**Fig 2:**
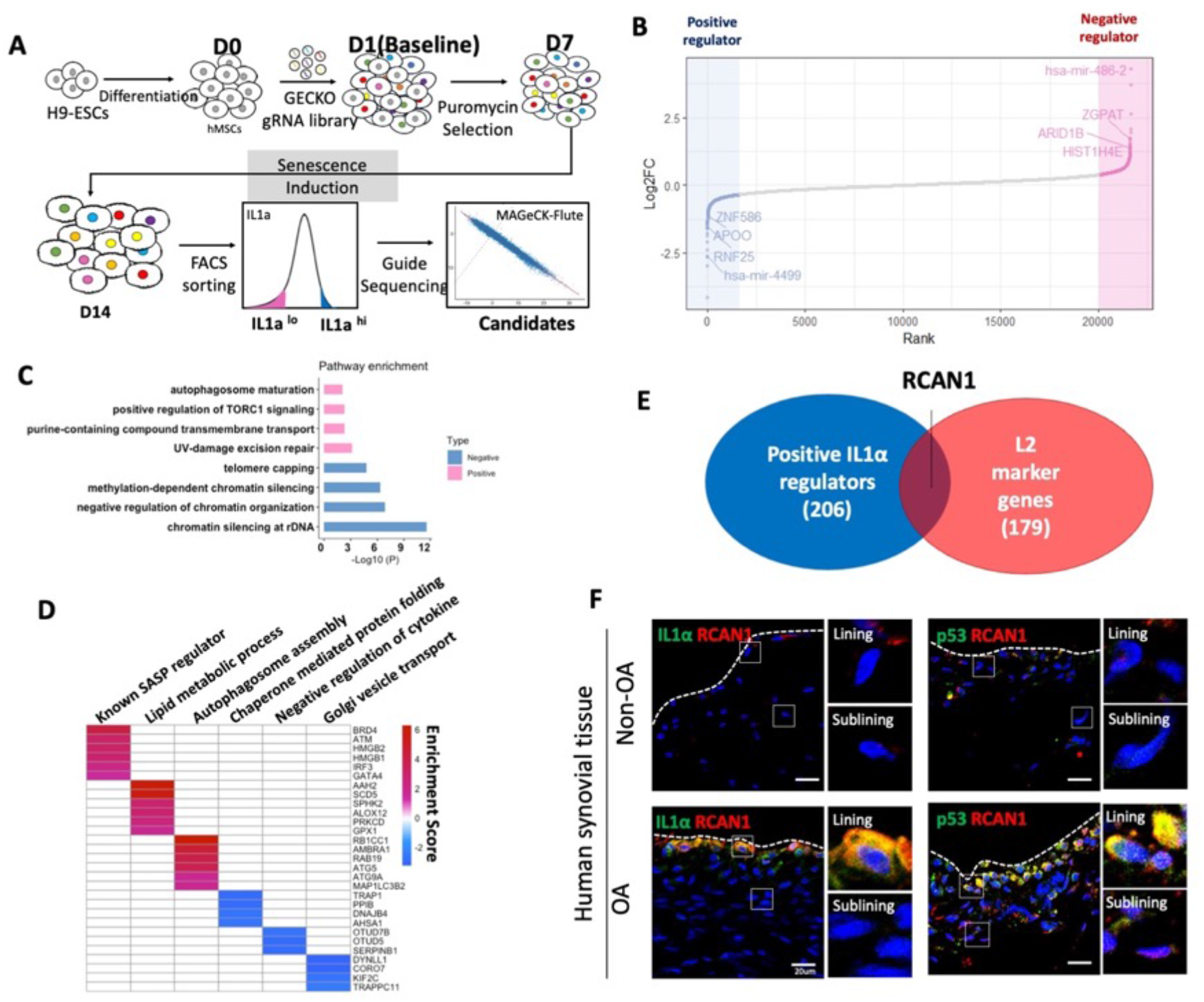
Genome-wide CRISPR screening identifies RCAN1 as candidate regulators of IL1α and co-expressed with IL1α in lining senescent cells. **A**, screen workflow. ESC-MSCs were infected with the library at a multiplicity of infection (MOI) of 0.3 and selected with puromycin for transduced cells. Cells were treated with doxorubicin for 7 days to establish cellular senescence and sorted based on high or low cell-surface IL1A expression. **B**, Rank-ordered normalized z-score (NormZ score). Hits at FDR < 5% are highlighted in blue (positive genes) and red (negative genes). Candidate genes were plotted based on mean log2 fold change of gRNA counts compared to control selection and p-values computed by MaGeCK. **C**, Pathway themes enriched in IL1A^low^ (positive) and IL1^high^ (negative) group. −log10(P) represents the −log10 of the adjusted P value. **D**, Selected pathways and corresponding genes identified in the screen. Color scale represents log2 fold change of average gRNA representation. **E**, Shared and unique genes identified as positive IL1A candidates in CRISPR screening and marker genes upregulated in L2 senescent lining cluster. **F**, Immunofluorescent staining of IL1α, p53 and RCAN1 in non-OA and OA synovium. Scale bar, 20μm.

### RCAN1 is a candidate regulator of IL1α and co-expressed with IL1α in lining senescent cells of OA synovium

To identify potential therapeutic target that modulate the senescence secretome of senescent fibroblasts in OA, we considered the candidate to be a hit as positive regulator of IL1α in CRISPR screening and an up-regulated marker in IL1α+ senescent fibroblasts (L2). Using these criteria, we validated ten candidates (Fig. 2E). Notably, Regulator of Calcineurin 1 (RCAN1) has been reported to play a pathogenic role in Down syndrome and Alzheimer’s disease^32,33^. We confirmed that RCAN1 was indeed co-expressed with p53 and IL1α in the human OA synovium (Fig. 2F). In confirm with this observation, Rcan1 positive cells also accumulated in cartilage, subchondral bone, and synovial tissue in mouse OA models (Fig. S2C). To determine the role of RCAN1 in cellular senescence, we used doxorubicin to induce senescence growth arrest in human ESC-MSCs. Notably, RCAN1 upregulation coincided with senescence growth arrest, SA-β-gal staining, and the upregulation of genes in the proinflammatory SASP wave such as IL1β, IL6 and IL8 (Fig. 3F-G, Fig. S2A-C). Collectively, these data suggest that the expansion of a potentially pathogenic population of synovial fibroblasts is marked by expression of RCAN1 and IL-1α.

**Fig 3:**
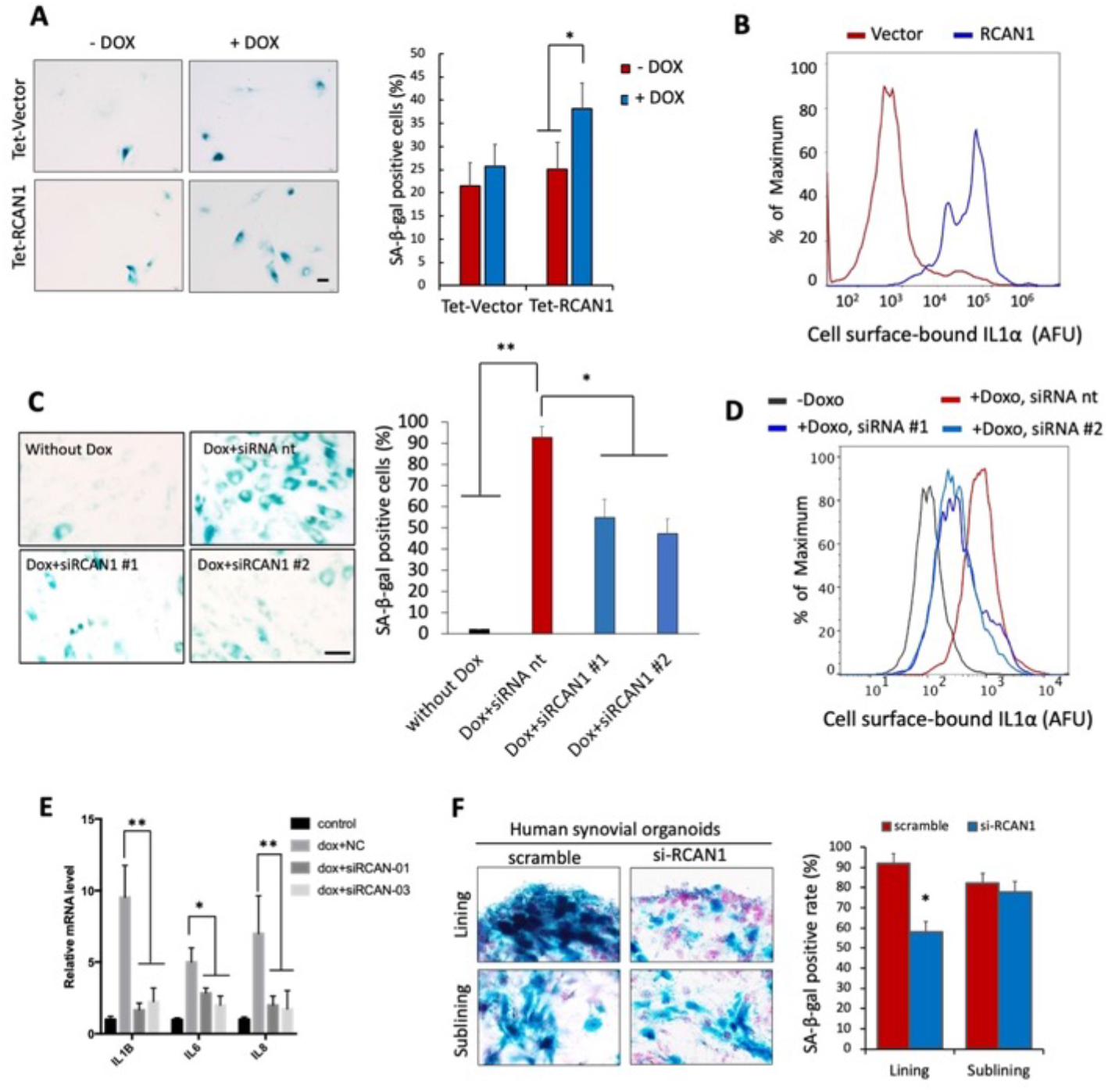
RCAN1 promotes the cellular senescence and proinflammation phenotype. **A-B**, Left: hESC-MSCs infected with either a Dox-inducible (Tet-On) vector expressing RCAN1 (Tet-RCAN1) or an empty vector (Tet-Vector) were grown with or without Dox for 9 days and SA-β-gal staining were analyzed (A), and cellular surface IL1α fluorescence was measured by fluorescence-activated cell sorter (FACS) (AFU, arbitrary fluorescence units) (B). Data are shown as percentage of maximum expression (i.e., the number of cells in each bin divided by the number of cells in the bin that contains the largest number of cells) for normalization. Scale bars, 50μm. **C-E**, hESC-MSCs was exposed to 0.2mM doxorubicin for 24h and transfected with indicated siRNAs every 3 days for 7 days, then the SA-β-gal staining were analyzed (C), cellular surface IL1α fluorescence was measured by FACS (D), and the abundance of mRNAs for the indicated proinflammatory SASP genes were quantified by qPCR (E). Scale bars, 50μm. **F**, Human synovial fibroblast organoids derived from OA patients were treated with indicated siRNAs every 3 days for 7 days, and SA-β-gal staining were analyzed and quantified. Scale bars, 20 μm. Statistical analysis: Two-way ANOVA was used for A, C, D and E. Two-sided Student’s t-test were used for B and F.Data represent the mean ± s.d for three independent experiments. *P < 0.05, **P < 0.01.

### RCAN1 promotes the cellular senescence and pro-inflammatory phenotype

To confirm the finding that RCAN1 regulate IL1A level in senescent cells from our CRISPR/Cas9 screening, we examined the effects of ectopic expression or depletion of RCAN1 during the senescence response of human ESC-MSCs. Ectopic expression of RCAN1 induced senescence in human ESC-MSCs, as shown by increased SA-b-Gal activity and cell-surface bounded IL1A level (Fig. 3A-B). Next, we treated human ESC-MSCs with doxorubicin, which causes a senescence growth arrest in more than 95% of cells and treated them with anti-RCAN1 or non-targeting siRNAs. Indeed, knockdown of RCAN1 via siRNA at the time of senescence initiation suppressed the cell surface-bounded IL1Α and expression of proinflammatory SASP (Figure 3C-D). Meanwhile, knockdown of RCAN1 decreased the expression of other pro-inflammatory SASP, including IL1β, IL6 and IL8 (Figure 3E). It is possible that the observed suppression of the SASP is an indirect effect of the suppression of senescence and its associated growth arrest through the inhibition of RCAN1. However, we established that knockdown of RCAN1, did not affect the growth and expression of cell-cycle arrest genes in the senescent cells. To investigate mimic the structure of human synovium in vivo, we used primary cultured human synovial organoid and cartilage tissue explants from patients undergoing total knee arthroplasty. Quantifications showed that anti-RCAN1 siRNA transfection but not non-targeting siRNA significantly suppress cellular senescence in lining layer of synovial organoids as evidenced by the induction of SA-β-Gal activity(Fig. 3F). Taken together, our results suggest that RCAN1 drives cellular senescence and pro-inflammatory SASP in senescent cells.

### RCAN1^+^IL1α^+^ senescent fibroblasts in the lining layer of synovium exhibit reprogrammed lipid metabolism

We next asked if our single cell RNA-seq dataset could provide novel insights into how RCAN1^+^IL1α^+^ senescent synovial fibroblasts contribute to OA pathogenesis. First, we examined the potential developmental relationship between the different clusters using the pseudotime algorithm monocle2^34^ on PRG4^high^ lining fibroblast subset and identified a senescent trajectory (Fig.4A). The expression of known SNC marker CDKN1A (p21^CIP1/WAF1^) along this trajectory suggests the L4 cluster as “the non-senescent root” and the RCAN1^+^IL1α^+^ cluster (L2) as “the senescent end” (Fig.4A-B). Displaying the expression of telomere maintenance-related genes, classic senescent and SASP marker genes, such as CCT5/7/8, CDKN1A, CCL2 and IL6, along the pseudotime also confirmed “the senescent end” annotation (Fig.4C). To further understand the process represented by the senescent trajectory, we identified genes that were dynamically regulated as the cells progressed through the trajectory and divided them into two classes: down-regulated (class A) and up-regulated (class B) along pseudotime (Fig.4D). Class A genes were enriched for the GO terms “telomere maintenance via telomerase”, “lysosome organization”, and class B genes were strongly enriched for “signal transduction by p53 class mediator” and “cellular senescence” (Fig.4D).

Mitochondrial dysfunction is a hallmark of senescence which causes metabolic reprogramming and distinct SASP^19,27,35^. It has been reported that increased RCAN1 level leads to mitochondria dysfunction^36-38^. Interestingly, GO terms associated with mitochondria function, including “ATP synthesis coupled electron transport” and “fatty acid catabolic process”, were the top regulated pathway that decreased along pseudotime (Fig.4D). Specifically, ACADVL, HADHB and other genes that encode mitochondrial enzymes that catalyze mitochondrial beta-oxidation of fatty acids, decreased along the pseudotime (Fig.4E). In contrast, the lipid biosynthesis genes, such as INSIG1 and FASN, were increased along the pseudotime (Fig.4E). The imbalance between lipid synthesis and consumption may be led to excessive accumulation of free lipids and lipotoxicity. Coincidently, “intrinsic apoptotic signaling pathway” and “response to oxidative stress” related genes were also increased along pseudotime (Fig.4D-E). Next, to evaluate whether RCAN1 expression was correlated with these meta-inflammatory phenotypes in RCAN1^+^IL1α^+^ cluster, we construct co-expression networks using single cell gene co-expression network analysis (scWGCNA)^39^ and robust modules of interconnected genes were identified (Fig.S3). Through analyze the distribution of module eigengenes in each of the lining synovial fibroblast clusters, we found that the brown module turned out to be specifically upregulated in the RCAN1^+^IL1α^+^ senescent cluster (L2) (Fig.4F). In addition, RCAN1 was identified as a hub gene of the brown module, and highly connected with genes such as SLC25A37 and BTG2 (Fig.4G). Finally, GO term enrichment analysis shows that major GO terms associated with the brown module were related to lipid metabolism, oxidative stress and cell cycle arrest (Fig.4H). Together, these results indicated that RCAN1^+^IL1α^+^ cluster represents the senescent end, which display mitochondrial dysfunction, imbalanced lipid metabolism, as well as increased lipotoxicity, which may be regulated by a network containing RCAN1.

**Fig 4:**
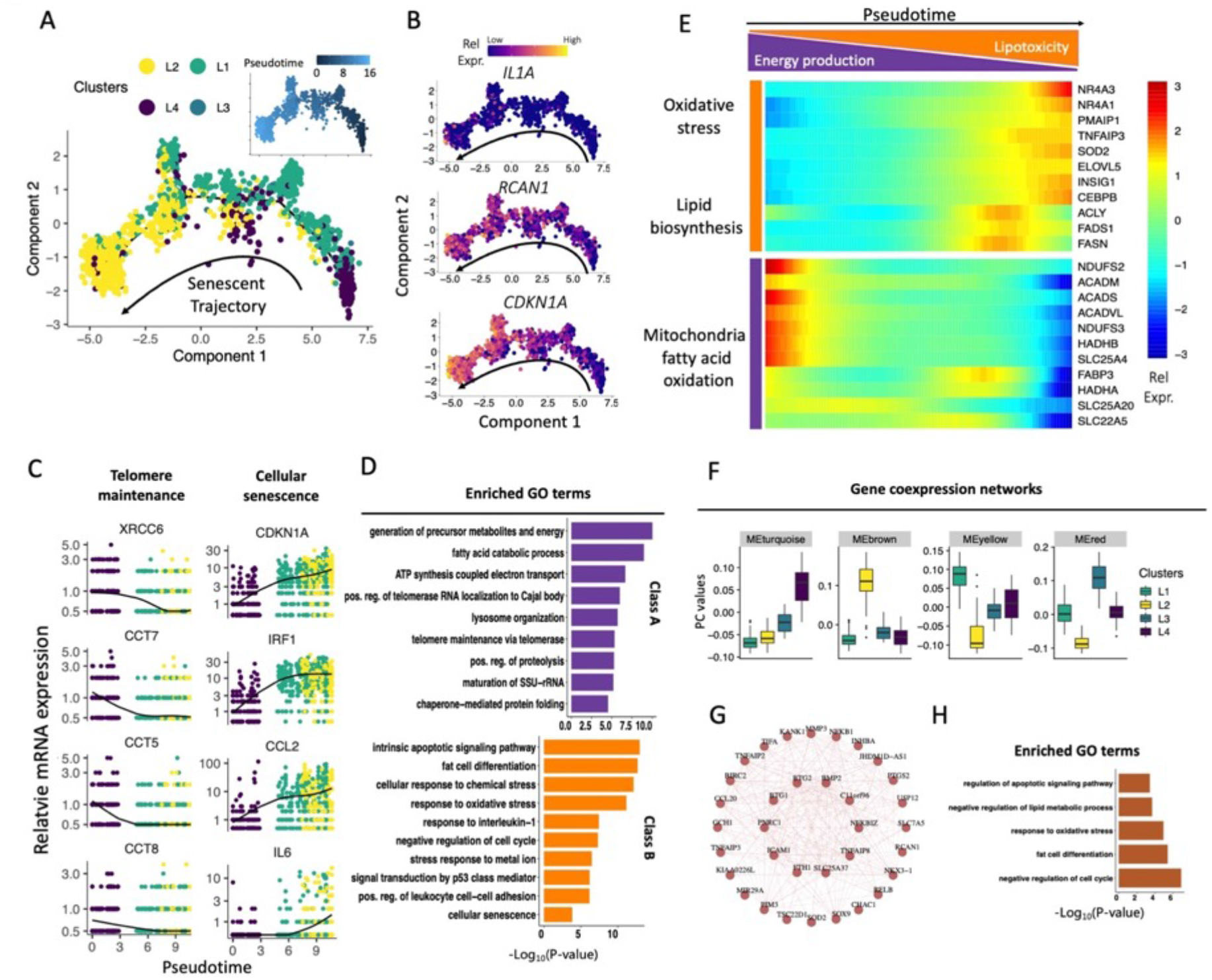
RCAN1^+^ IL1α^+^ senescent fibroblast in the lining layer of synovium shows enhanced lipid metabolism and lipotoxicity. **A**, Monocle pseudospace trajectory revealing the OA lining fibroblasts senescent progression colored according to cell clusters. **B**, Cell trajectory projections of transcriptional changes in CDKN1A, RCAN1 and IL1A genes based on the manifold. **C**, Pseudotime projections of transcriptional changes in senescence relevant genes based on the manifold. Each dot represents a cell and colored according to cell clusters, the solid line represents the loess regression. **D**, GO terms from enrichment analysis reveal the different function of the senescent and non-senescent lining fibroblasts. **E**, Heatmap showing the gene expression dynamics of the OA lining fibroblasts along the senescent trajectory. Genes were grouped by similar expression profiles with hierarchical clustering. **F**, Results of scWGCNA. Eigengene expression of four selected modules across all 4 lining fibroblast clusters. Color code of the modules is preserved. The brown module has enriched expression in cluster L2 (RCAN1+ IL1α+ senescent fibroblasts). **G**, Hub-gene network of the brown module (the module containing RCAN1). **H**, Eigengene expression of module brown were subjected to GO terms enrichment analysis.

### Saturated fatty acid secretion from senescent synovial fibroblasts promotes cartilage degeneration and could be inhibited by si-RCAN1

To further evaluate the role of RCAN1 in abnormal lipid metabolism in senescent cells, we performed RNA sequencing from mouse fibroblasts transfected with si-Ctrl and si-Rcan1. Unsupervised cluster analysis segregated si-Ctrl from si-Rcan1 group and revealed prominent differences between their transcriptome, with 692 significantly differentially expressed genes (Supplementary table1). Pathway analysis indicated that the identified gene sets are associated with annotations related to “Aging” and “cellular response to interleukin-1” (Figure 5A). Interestingly, “Response to fatty acid” was significantly downregulated and “sphingolipid metabolic process” was activated after RCAN1 inhibition, indicating reduced level of intracellular free fatty acid and increased channeling of fatty acid into structural lipids, such as sphingolipids (Figure 5A). Specifically, *de novo* fatty acid synthesis genes, such as ACLY, FASN and ELVOLs was downregulated, while genes involved in channeling free fatty acid to mitochondria oxidation, peroxisome and membrane structure were upregulated upon RCAN1 knockdown (Figure 5B).

**Fig 5:**
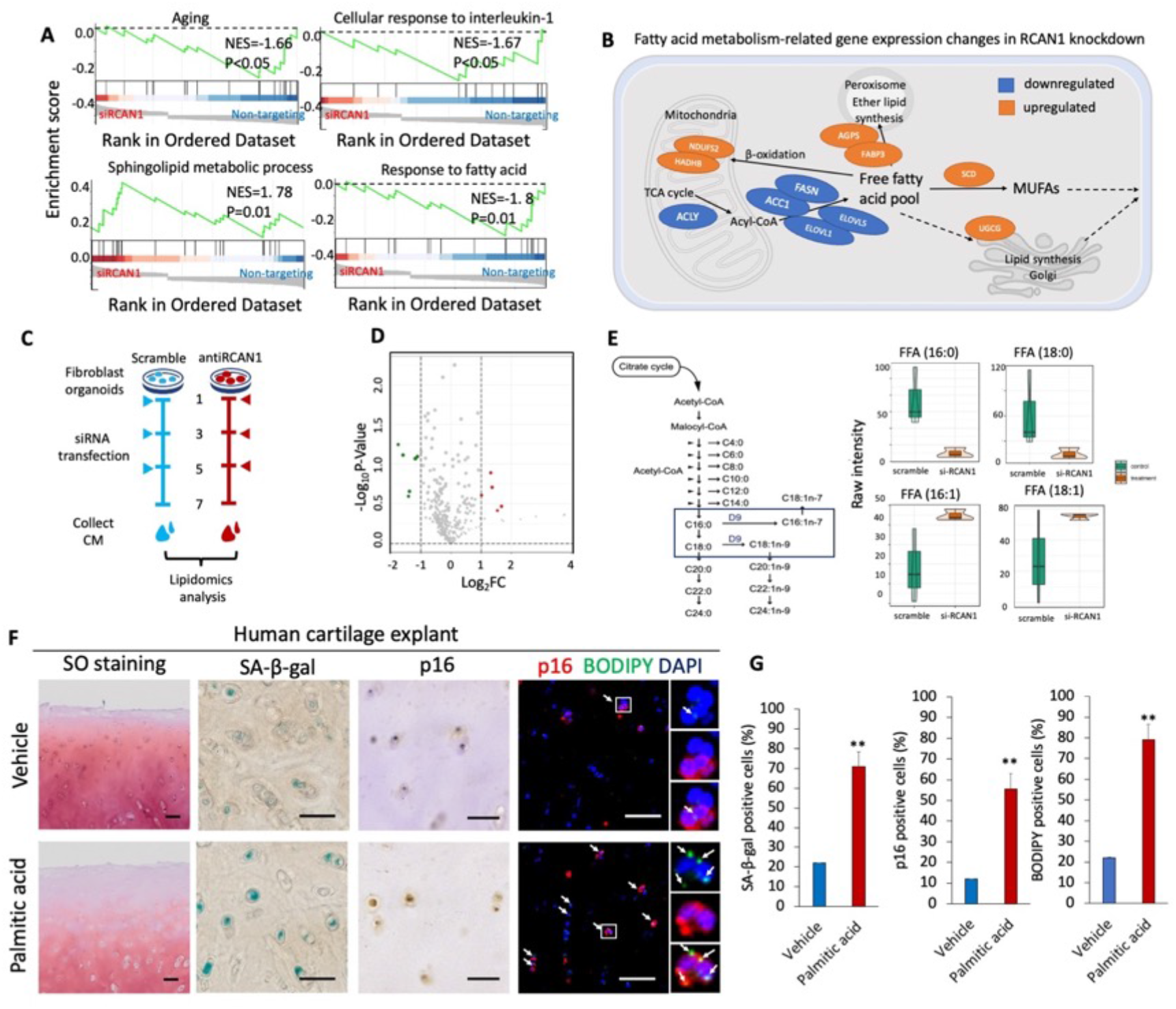
Saturated fatty acid secretion from senescent synovial fibroblasts promotes cartilage degeneration and could be inhibited by si-RCAN1. **A**, mRNA expression profiling from senescent fibroblasts treated with non-targeting (si-Ctrl) or anti-Rcan1 (si-Rcan1) were analyzed with RNA sequencing. Gene Set Enrichment Analysis (GSEA) showing enrichment of pathways between si-Ctrl and si-Rcan1. **B**, Pathway map of genes related to fatty acid metabolism that are differentially expressed in between si-Ctrl and si-Rcan1. **C**, Experimental schematic illustration. Conditioned medium (CM) from synovial organoids treated with indicated siRNA were collected and lipidomics analyzed with UPLC-MS/MS. **D**, Volcano plot shows differentially detected compounds in lipidomics analysis of CM from senescent synovial organoids treated with or without anti-RCAN1 siRNA (n = 3). **E**, Left: Fatty acid synthesis pathway. Right: Violin plots shows saturated and monounsaturated fatty acid levels. **F-G**, Human cartilage explants obtained from patients undergoing total knee arthroplasty were cultured ex vivo and treated with vehicle (DMSO) or 10 uM palmitic acid for 7 days (n=3). Sections were stained with SO, SA-β-gal, p16 and BODIPY (lipid droplets) (F) and quantified (G). White arrows indicate lipid droplets. Scale bar, 50μm.

SASP has been characterized largely for secreted proteins and the lipid components of the SASP are understudied, but recent study suggests that bioactive lipids such as prostaglandins, fatty acids and ceramides also comprise an important part of SASP^19,20^. To explored if and how the fatty acid metabolism was related to senescent synovial fibroblasts induced cartilage degeneration, we used primary cultured human synovial organoid and cartilage tissue explants from patients undergoing total knee arthroplasty. To identify the secreted lipid factors regulated by RCAN1, we compared the lipidomics analysis of conditioned medium from senescent synovial organoids treated with or without anti-RCAN1 siRNA (Figure 5C). We identified 11 differential expressed lipids that secreted from senescent synovial fibroblasts (Figure 5D). Notably, two of mono-unsaturated free fatty acid, palmitoleic acid (FFA (16:1)) and Oleic acid (FFA (18:1)), were up-regulated in anti-RCAN1 siRNA treated group, while two saturated FFA, palmitic acid (FFA (16:0)) and Stearic acid (FFA (18:0)) are down-regulated (Fig. 5E). Accordingly, SCD, a gene that encode a desaturase which introduce the first double bond into saturated fatty acyl-CoA substrates and gives rise to a mixture of 16:1 and 18:1 unsaturated fatty acid, was up-regulated in in anti-RCAN1 siRNA treated group (Figure 5B). To functionally test whether these senescent synovial fibroblast derived lipid mediators are involved in cartilage degeneration, we used in vitro cartilage tissue explants. We found that palmitic acid induced cellular senescence, lipid droplet formation and extracellular degeneration in human cartilage tissue explants (Fig. 5F-G). Thus, these results suggest that RCAN1 may regulates pro-degenerative paracrine effect of senescent synovial fibroblasts through saturated fatty acids, such as palmitic acid and Stearic acid.

### Silencing of Rcan1 in the lining synovial fibroblasts counteracts OA progression *in vivo*

The therapeutic effect of Rcan1 inhibition was examined in destabilization of the medial meniscus (DMM) induced OA model in mice (Fig. 6A). Intra-articular delivery of siRNA labeled with Cy3 indicated uptake of the siRNA by lining layer fibroblasts in synovial tissue (Fig. 6B). Administration of si-Rcan1 efficiently inhibited Rcan1 expression in cartilage, subchondral bone, and synovial tissue (Fig. 6C). Severe cartilage destruction, subchondral bone sclerosis, osteophyte development, and a moderate degree of synovial inflammation was observed in si-Ctrl group (Fig. 6D-E). While, intra-articular injection of si-Rcan1 ameliorated DMM-induced cartilage destruction and subchondral bone sclerosis, along with evident effects on synovial inflammation (Fig. 6D-E). Specifically, Rcan1 inhibition in joint synovium of OA mice caused concomitant recovery of col2a synthesis and suppression of both inflammatory SASP factors and non–cell autonomous propagation of cellular senescence in cartilage (Fig. 6F-G). Moreover, we measured lipid accumulation in OA joints si-Ctrl and si-Rcan1 treated legs. DMM surgery with intra-articular injection of si-Ctrl caused accumulation in the OA joint, which was alleviated by treatment with si-Rcan1(Fig. S4). Together, this indicates that Rcan1 antagonism effectively ameliorates OA progression in mice.

**Fig 6:**
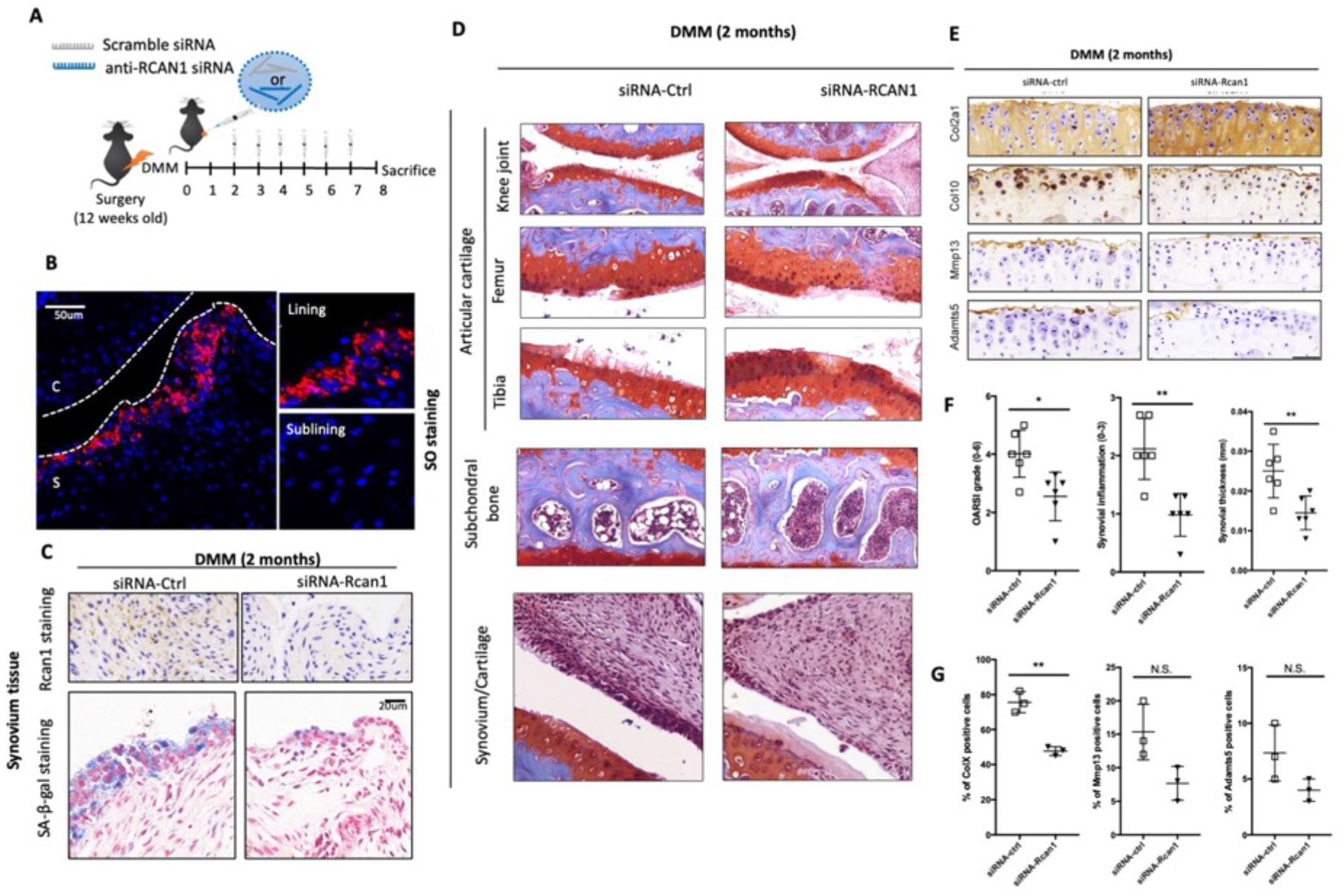
Silencing of Rcan1 in the lining synovium counteracts OA progression *in vivo*. **A**, Schematic illustration of si-RNA therapy in posttraumatic OA model mice. Anti-Ctrl or anti–Rcan1 siRNA was intra-articularly delivered to sham- or DMM-operated mice using an in vivo delivery system, MSN-PEI nanoparticles (0.4 mg/kg; once every 7 days for 6 weeks). **B**, Confocal imaging of distribution of si-RNA labeled with Cy3 in treated mouse joints. **C**, SA-β-gal staining and immunohistochemistry of Rcan1 were conducted in synovial sections from mice after intra-articular delivery of siR-Ctrl or siR-Rcan1. Scale bar, 20μm. **D**, Joint sections were stained with safranin O. The inset in the images is shown at magnified images in the bottom row. Scale bar, 200μm. **E**, Cartilage destruction, subchondral bone sclerosis, osteophyte formation, and synovial inflammation were determined by safranin O/hematoxylin staining and scored (n = 6). **F-G**, Extracellular matrix (Col2, Col10a), SASP factors (Mmp13 and Adamts5) were detected by immunohistochemistry (IHC) (n = 3). Scale bar, 25μm.

### Silencing of RCAN1 in synovial tissue effectively attenuates chondrocyte senescence in human OA cartilage

Last, we evaluated the relevance of our findings to humans using primary cultured human articular tissue explants from patients undergoing total knee arthroplasty (Fig.7A). Anti-RCAN1 siRNA transfection but not non-targeting siRNA significantly suppress RCAN1 and IL1A expression level, as well as cellular senescence (characterized by SA-β-Gal staining) in lining layer of human OA synovial tissue explants (Fig. 7B). We than cocultured the OA cartilage explants (from un-affected cartilage area) with synovium explants which has been pre-treated with anti-RCAN1 siRNA or non-targeting siRNA (Fig.7A). Quantifications showed that chondrocytes senescence and matrix degeneration in human cartilage explants coculture with anti-RCAN1 siRNA treated synovium was inhibited compared with non-targeting siRNA treated synovium as evidenced by the SO staining, Col2a1, p16 and SA-β-Gal staining (Fig. 5C-E). Taken together, we found that inhibition of RCAN1 in synovial tissue could suppress cellular senescence and matrix degeneration in cartilage tissue.

**Fig 7:**
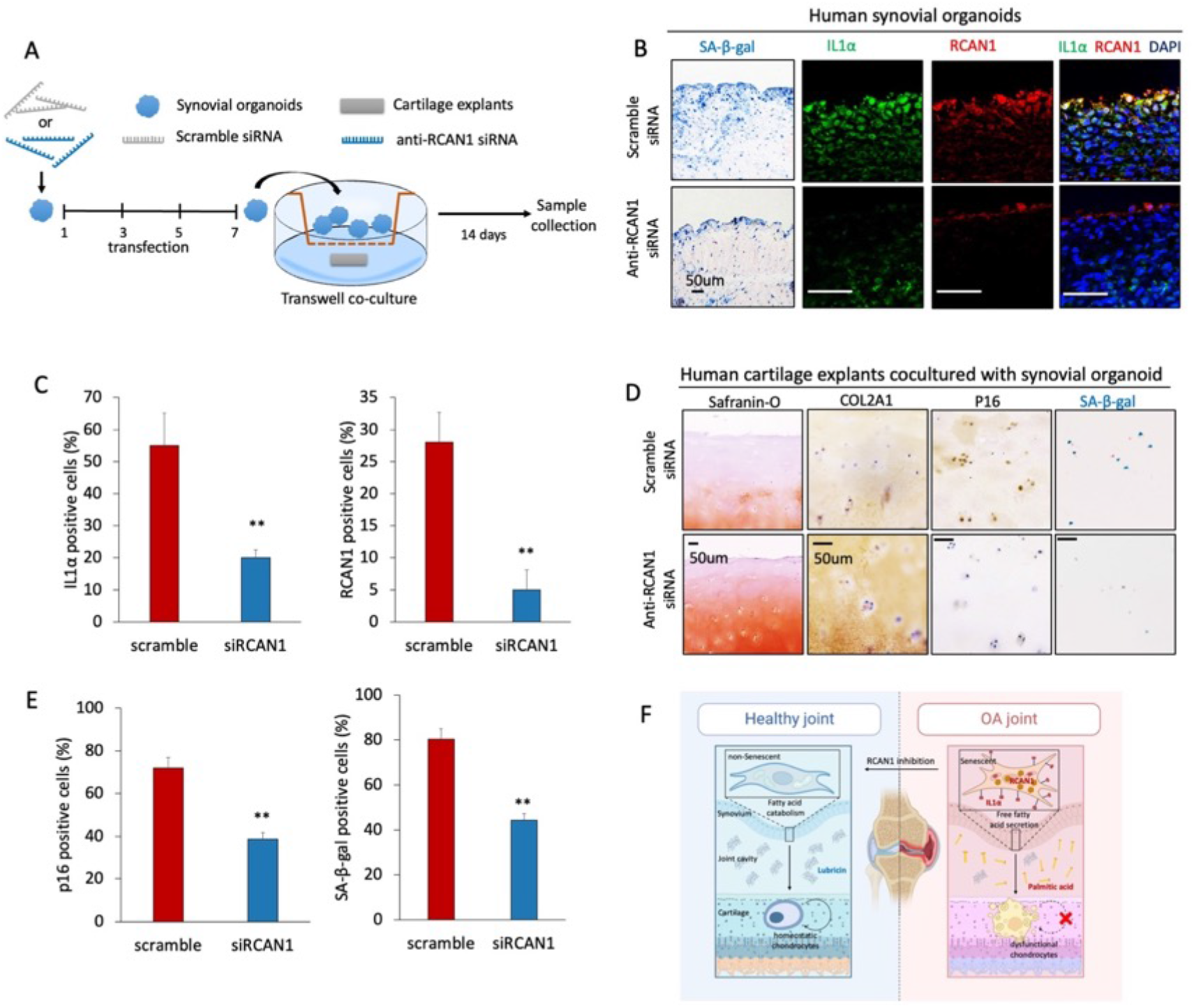
Silencing of RCAN1 in synovial tissue effectively attenuates chondrocyte senescence in human OA cartilage. **A**, Experimental schematic illustration. Synovial organoids derived from and OA synovial fibroblasts were treated either with scramble siRNA or anti-RCAN1 siRNA every 2 days for 7 days and co-cultured with human OA cartilage explants for another 14 days. B-C, Synovial organoids sections treated either with scramble siRNA or anti-RCAN1 siRNA were stained with IL1α and RCAN1 (B) and the percentage of cells positive for IL1α and RCAN1 were quantified (n ≥ 3) (C). Scale bar, 50μm. D-E, Cartilage explant sections were stained with safranin O/hematoxylin, SA-β-gal/Nuclear Fast Red, p16 and COL2A1 (D). The percentage of cells positive for SA-β-gal, p16 and COL2A1 was quantified (n ≥ 3) (E). Scale bar, 50μm. F, Graphic summary of this study: RCAN1 in a IL1α+ subsets of synovial fibroblasts accelerate cartilage degeneration via saturated free fatty acids (FFAs).

These findings suggest that increased cellular senescence and matrix degeneration in cartilage observed in OA patients may in part be triggered by impairment in lipid homeostasis in synovial fibroblasts.

## DISCUSSION

Osteoarthritis (OA), an age-related degenerative disease, is closely associated with progressive accumulation of senescent cells (SNCs) in cartilage, which has a causative role of cartilage destruction. Although the presence of SNCs in OA synovium has been noted, little is known about SNCs heterogeneity or how SNCs subset in the synovium affect the whole-joint environment and OA development. Here, combining single-cell RNA sequencing (scRNA-seq) and genome wide CRISPR/Cas9 screening, we identify a novel regulator of SASP, RCAN1, in a distinct IL1Α positive subset of senescent synovial fibroblasts which is responsible for mediating chronic synovial inflammation and OA progression. And we demonstrate that targeting RCAN1 with siRNA, is a disease-modifying agent in OA that prevents cartilage degeneration and promotes matrix regeneration (Fig. 7F).

Our study suggests a possible alternative approach to the recently developed senolytic therapy for OA treatment. Cellular senescence has recently emerged as a fundamental aging mechanism that contributes to OA development^7^. Elimination of senescent cells with senolytics UBX0101 showed promise as a potential disease-modifying therapy for osteoarthritis in mice, but a phase 2 clinical trial (NCT04349956) with the same drug failed to achieve primary endpoint of improving pain in patients with knee osteoarthritis, requiring the development of alternative therapeutic strategies that interfere with the detrimental effects of senescent cells in OA. Because chondrocytes are sparse in the human cartilage, excessive reduction of the chondrocyte population may accelerate exhaustion of regenerative cell populations that potentially replace chondrocytes during OA process and contribute to cartilage repair^7^. Another important consideration is that removing senescent cells can lead to unwanted outcomes, which could affect not only non-senescent cells but also beneficial senescent cells, such as those participating in wound repair ^40-42^. Since many of the negative effects associated with senescence in osteoarthritis are driven by the SASP, compounds that modulating senescence secretome, often referred to as senomorphics, may serve as an attractive strategy to treat osteoarthritis^5,8,14^. In our study, we found that a subset of IL1Α+p16+ senescent cells was expanded in human OA synovium compared with normal synovium, which was also observed in a mouse model of ACLT-induced OA mouse^7^. These senescent synovial fibroblasts upregulate catabolic SASP factors (e.g., MMP3, PTSG2, PLAUR, and SERPINE1) and contribute to cartilage destruction. Furthermore, due to the distinct superficial location at loose synovium tissue, efficient gene silencing in these senescent cells can be easily achieved while circumventing excessive or unintended removal of cell populations in cartilage. Together, we reported that siRNA therapy targeting RCAN1 in senescent synovial fibroblasts may effectively quenching of pro-degenerative SASP factors while enable sustained matrix anabolism by chondrocytes.

High-throughput siRNA- or shRNA-based screening, pooled CRISPR-Cas9–based knockout screening has enabled reliable systematic identification of genes controlling cellular senescence in humans and allowed the discovery of targets for aging interventions ^43-45^. These studies mainly focused on the regulation pathway of cell survival and cell cycle arrest in senescent cells as they use cell proliferation as the readout of the screenings ^43-45^. Although SASP mediated many detrimental effects of senescence through paracrine action, systematic dissect of SASP regulatory pathways is still lacking. Here, we used the CRISPR-Cas9-based gene knockout strategy, a pooled library of sgRNAs covering the entire annotated human genome, and human stem cell models of DNA damage induced senescence to enable the systematic identification of IL1Α regulatory genes on a genome-wide scale. Consistent with previous studies, we also identified a panel of known IL1Α and SASP regulators such as BRD4, GATA4, HMGB2 and ATM^17,31,46-49^. In addition, we uncovered several previous unknown candidate genes involved in lipid metabolic process, chaperone mediated protein folding pathways which may participated in SASP regulation during senescence. Thus, our effort expanded our understanding of human SASP regulatory factors and potential therapeutic targets.

Our genome-wide screen and functional validation showed that the regulator of Calcineurin 1 (RCAN1) governs the proinflammatory and pro-degenerative senescence-associated secretome, as loss of this gene resulted in decreased IL1Α level and delayed cellular senescence. Although RCAN1 has been previously reported as a player in pathogenesis of Down syndrome and Alzheimer’s^32,33,36,50,51^, our study adds a layer of complexity to RCAN1 function by revealing its important role in SASP regulation and pinpointing RCAN1 as a new therapeutic target for treating aging-associated osteoarthritis. In addition to a disease-modifying effect, the absence of side effects also makes RCAN1 inhibition stand out compared to other senomorphics interventions such as rapamycin. Mammalian target of rapamycin (mTOR) is a key post-transcriptional regulator of pro-inflammatory and pro-tumorigenic SASP in many cellular senescence and disease settings^17,31,52^. mTOR inhibition with rapamycin protects mice from osteoarthritis^53^, but rapamycin, also used as an anticancer drug, has severe side effects. In our study, we treated mice with siRcan1 for 8 weeks without observing any obvious side effects or toxicity. Along the same line, the siRNA technology we used to silencing RCAN1 signaling in vivo is already used in various clinical settings^54,55^.

Our findings also support a link between lipid metabolism and OA development. OA is a complex disease complicated with several metabolic diseases including dyslipidemia, hyperglycemia, and hypertension^3,4^. Several metabolites, such as total free fatty acid (FFA) have been reported to be increased in OA patients^56,57^. Notably, several studies revealed that lipid metabolic process such as cholesterol metabolism and de novo lipogenesis in chondrocytes are a crucial catabolic regulator of the pathogenesis of osteoarthritis^22,23^. Although the SASP has been characterized largely for secreted proteins, the lipid components of the SASP have also been recognized recently^19,20^. However, whether a metabolite itself is a causal factor in OA progression remain elusive. Our results show that saturated fatty acids are lipid component of the SASP secreted by senescent synovial fibroblasts and consequently promotes the cellular senescence and matrix degeneration in chondrocytes.

Potential limitations of our study include the use of intra-articular injection–mediated delivery of Rcan1 rather than the use of transgenic mice with a fibroblast-specific promoter for Rcan1 knockout. Although we confirmed that synovial lining fibroblasts are the target of transfection by the jetPEI and miRNA complex, the delivery of the injected siRNA to the chondrocyte cannot be ruled out. The transgenic knockout of Rcan1 under the Prg4 promoter would more clearly dissect the role of Rcan1 in synovial fibroblasts in mediating OA development in mice. Another limitation reflects the technical hurdles is that a treatment regimen of weekly intra-articular injections is unfeasible in patients. Therefore, development of protective vehicles or RNA modification methods will be necessary to design a therapeutic strategy with a practical injection interval for anti-RCAN1 siRNA. Last, we validated RCAN1 inhibition in a model of instability-induced osteoarthritis. An approach to decrease RCAN1 activity needs to be further tested in large animals similar in anatomy and joint mechanics to humans. Whether RCAN1 inhibition would be effective in patients whose osteoarthritis is driven by different mechanisms—obesity or genetic factors—is still to be tested.

## MATERIALS AND METHODS

### Study design

The overall objective of this study was to investigated how senescent synovial fibroblasts contribute to OA pathogenesis and assessed whether the joint homeostasis can be restored by novel anti-senescence approaches. This study included in vivo and in vitro experiments using samples from patients with end-stage osteoarthritis or C57BL/6 mice that had undergone a surgical osteoarthritis models. Primary chondrocytes and synovial fibroblasts from patients and C57BL/6 mice, or H9 human mesenchymal stem cells derived from embryonic stem cells (hESC-MSCs) were used for in vitro experiments. We conducted single cell RNA sequencing of synovial tissues from non-OA patients (n=3) and OA patients (n=2). After quality control and integration with publised data (GSE152805), we obtained samples up to n=3 for non-OA patients and n=3 for OA patients. For in vitro human tissue explants culture, human OA cartilage and synovium specimens were collected from patients with OA underwent total knee replacement at Zhejiang Provincial People’s Hospital. The Institutional Review Board (IRB) of Zhejiang Provincial People’s Hospital approved the collection and use of these materials(IRB NO. 2019KY072). Full written informed consent was provided by all participants before the total knee replacement arthroplasty operative procedure. Mice used for animal studies were randomly assigned to each group and sample size reflects the number of independent biological replicates and is indicated in the figure legend. All animal experiments were approved by by the Zhejiang University Ethics Committee (NO. ZJU20190780). The design, analysis, and reporting of animal experiments followed the Animal Research: Reporting of In Vivo Experiments guidelines. In the genome wide CRISPR/Cas9 screening, the human GeCKOv2A pooled knockout library was used to identify genes responsible for IL1α expression in senescent hESC-MSCs cells. The library was a gift from Feng Zhang lab (Addgene # 1000000049)^58^. The sgRNA read count and hits calling were analyzed by MAGeCK v0.5.7 algorithm^59^.

### Statistical analysis

For in vitro studies, each experiment was conducted independently at least three times. To compare the experimental groups, parametric test based on unpaired two-tailed Student’s t test or one-way ANOVA was used. For in vivo experiments, each independent trial was conducted using an individual mouse. Previous experiments have been conducted under similar conditions in our laboratory (same model but different drugs) to estimate a biologically relevant effect size and SD. To determine significant differences, nonparametric test based on Mann-Whitney U test was used. For nonparametric, multigroup comparisons, the Mann-Whitney U test was used. Values of P < 0.05 were considered statistically significant. Significance level was presented as either *P < 0.05 or **P < 0.01.

## Acknowledgements

We would like to acknowledge the technical support by the Core Facilities, Zhejiang University School of Medicine. We also thank all group members of H.W. lab for helpful discussions.

## Funding

This study was supported by the National key R&D program of China (2017YFA0104900), the National Natural Science Foundation of China (NO. T2121004, 31830029, 81802146 and 81802195).

## Author contributions

Conceptualization: O.H., and H.J.; methodology: H.J., Y.D., W.D., and W.J.; formal analysis: H.J., Y.D., and W.D.; collected and inspected human patient samples: Y.D., and C.Y.; single cell RNA sequencing and analysis: H.J., and W.J.; mouse DMM model and treatment: H.J., D.X., and W.D.; writing original draft: H.J., and W.J.; writing-review and editing: J.J., and O.H.; All authors read and edited the manuscript. O.H. supervised the study.

## Data availability

The raw sequence data reported in this paper have been deposited in the Genome Sequence Archive (Genomics, Proteomics & Bioinformatics 2021) in National Genomics Data Center (Nucleic Acids Res 2022), China National Center for Bioinformation / Beijing Institute of Genomics, Chinese Academy of Sciences (GSA-Human: HRA003571) that are publicly accessible at https://ngdc.cncb.ac.cn/gsa-human. The full genome wide CRISPR screening data will be available upon requests to the corresponding authors.

## Notes

### Competing Interest Statement

A patent application (202111459686.7; filed on 12/02/2021) has been submitted based in part on results presented in this manuscript on the use of siRNAs that target RCAN1 for osteoarthritis treatment. Hongwei Ouyang, Jiajie Hu and Dongsheng Yu are listed as the inventors.

